# Construction of soil defined media using quantitative exometabolomic analysis of soil metabolites

**DOI:** 10.1101/151282

**Authors:** Stefan Jenkins, Tami L. Swenson, Rebecca Lau, Andrea Rocha, Alex Aaring, Terry C. Hazen, Romy Chakraborty, Trent Northen

## Abstract

Exometabolomics enables analysis of metabolite utilization of low molecular weight organic substances by soil isolates. Environmentally-based defined media are needed to examine ecologically relevant patterns of substrate utilization. Here, we describe an approach for the construction of defined media using untargeted characterization of water soluble soil metabolites. To broadly characterize soil metabolites, both liquid chromatography mass spectrometry (LC/MS) and gas chromatography mass spectrometry (GC/MS) were used. With this approach, 96 metabolites were identified, including amino acids, amino acid derivatives, sugars, sugar alcohols, mono- and di-carboxylic acids, osmolytes, nucleobases, and nucleosides. From this pool of metabolites, 25 were quantified. Water soluble organic carbon was fractionated by molecular weight and measured to determine the fraction of carbon accounted for by the quantified metabolites. This revealed that, community structures, these soil metabolites have an uneven quantitative distribution, with a single metabolite, trehalose accounting for 9.9 percent of much like soil microbial the (< 1 kDa) water extractable organic carbon. This quantitative information was used to formulate two soil defined media (SDM), one containing 23 metabolites (SDM1) and one containing 46 (SDM2). To evaluate SDM for supporting the growth of bacteria found at this field site, we examined the growth of 30 phylogenetically diverse soil isolates obtained using standard R2A medium. The simpler SDM1 supported the growth of up to 13 isolates while the more complex SDM2 supported up to 25 isolates. One isolate, *Pseudomonas corrugata* strain FW300-N2E2 was selected for a time-series exometabolomics analysis to investigate SDM1 substrate preferences. Interestingly, it was found that this organism preferred lower-abundance substrates such as guanine, glycine, proline and arginine and glucose and did not utilize the more abundant substrates maltose, mannitol, trehalose and uridine. These results demonstrate the viability and utility of using exometabolomics to construct a tractable environmentally relevant media. We anticipate that this approach can be expanded to other environments to enhance isolation and characterization of diverse microbial communities.

**Highlights:**

- LC/MS and GC/MS analyses of soil extracts revealed a diversity of 96 metabolites.
- Soil defined media were constructed based on water extractable soil metabolomics data.
- The defined media supported the growth of 25 out of 30 bacterial isolates.
- Exometabolomics demonstrated preferential consumption of amino acids for one isolate.
- These media can be used to understand environmentally relevant microbial substrate preferences.

**Abbreviations:** DOM dissolved organic matter; SOM soil organic matter; WEOC water extractable organic carbon; LMWOS low molecular weight organic substances; SDM(1/2) soil defined media (1 and 2); R2A Reasoner’s 2A agar medium; ORFRC Oak Ridge Field Research Center; LC/MS liquid chromatography mass spectrometry; GC/MS gas chromatography/ mass spectrometry; TOC total organic carbon; HILIC hydrophilic interaction liquid chromatography

## 1. Introduction

Soil organic matter, historically considered to be composed of large polymeric humic substances, is now thought to largely consist of microbial products (Schmidt et al., 2011). Traditionally, the water soluble fraction of soil carbon, known as dissolved organic carbon or matter (DOC/DOM), is defined as the fraction that passes through a 0.45 μm filter (Ohno et al., 2014). Many microbes process macromolecular substrates such as plant biomass extracellularly and uptake the resulting low molecular weight organic substances (LMWOS). Thus, it is not surprising that the water extractable organic carbon (WEOC)(Boyer and Groffman, 1996; Guigue et al., 2014) is associated with high microbial activity and soil respiration (Haney et al., 2012). Examination of the composition of LMWOS using metabolomic methods has revealed a diversity of small molecule metabolites (Warren, 2014; Swenson et al., 2015b). Recently, we observed exometabolite niche partitioning in sympatric soil microbes indicating that there can be a strong linkage between the composition of soil exometabolites and microbial community structure (Baran et al., 2015). This reinforces long-standing views that it is desirable for culture media to approximate the conditions, especially the quality and abundance of metabolites found within their habitat. This raises the exciting possibility that soil metabolomics methods can help inform the development of relevant culture media to enable laboratory studies investigating microbial resource partitioning.

Most of our understanding of the biology of soil microbes is based on the small fraction of microbes that have been successfully cultivated in isolation. Reasoner’s 2A agar medium (R2A) is one of the most widely used for isolations and was developed for the cultivation of bacteria found in potable water (Reasoner and Geldreich, 1985). Its effectiveness may be attributable to its rich nutrients (peptone, casamino acids and yeast extract). However, the composition of R2A is dramatically different from the composition of soil LMWOS, and therefore has limited ecological relevance. There are many promising new technologies for isolating soil microbes with direct connection to the native small molecule environment (Pham and Kim, 2012) such as the use of transwell plates, soil substrate membranes, and recently the iCHIP that enables diffusion of native DOM into the plates for isolations (Svenning et al., 2003; Ferrari et al., 2008; Ling et al., 2015).

Exometabolomics enables the study of the transformation of the small molecule environment by bacteria and other microorganisms by comparing spent media to non-inoculated controls, typically using mass spectrometry. This direct assessment of phenotype provides high-resolution of metabolite depletion which is emerging as a powerful complement to existing broader techniques such as bulk soil respiration and carbon transformation measurements (Butler et al., 2003; Miller et al., 2005; Schimel and Mikan, 2005; Fernandes et al., 2013; Tucker et al., 2014). Previous studies have demonstrated the capability of exometabolomics in providing a functional complement to genomic and transcriptomic data (Baran et al., 2013; Liebeke and Lalk, 2014). Recently, exometabolomics was used with soil microbes and a microbial extract media to discover exometabolite niche partitioning for the 13 isolates and 2 environments examined (Baran et al., 2015). However, a limitation of this approach is that many of the metabolites in these environmentally relevant rich media could not be identified and presumably many others were not detected. For many exometabolite profiling experiments it would be desirable to have an environmentally relevant defined medium such that all of the metabolites are accounted for and the observed patterns of metabolite utilization are relevant to the field site.

Here we describe the construction of defined media for exometabolomics experiments based on untargeted analysis of metabolites from a specific soil sample of interest (Figure 1). Water extractable metabolites, referred to here as WEOC, were first qualitatively characterized using both liquid chromatography mass spectrometry (LC/MS) and gas chromatography mass spectrometry (GC/MS). From these data, a subset of metabolites was selected for absolute quantification to assist in formulating defined media that approximates the quality and quantities of metabolites within the soil. We evaluated the ability of the resulting media to support the growth of 30 phylogenetically diverse isolates from the Oak Ridge Field Research Center (ORFRC) and performed time series characterization of the substrate preferences of a single isolate.

**Figure 1.**
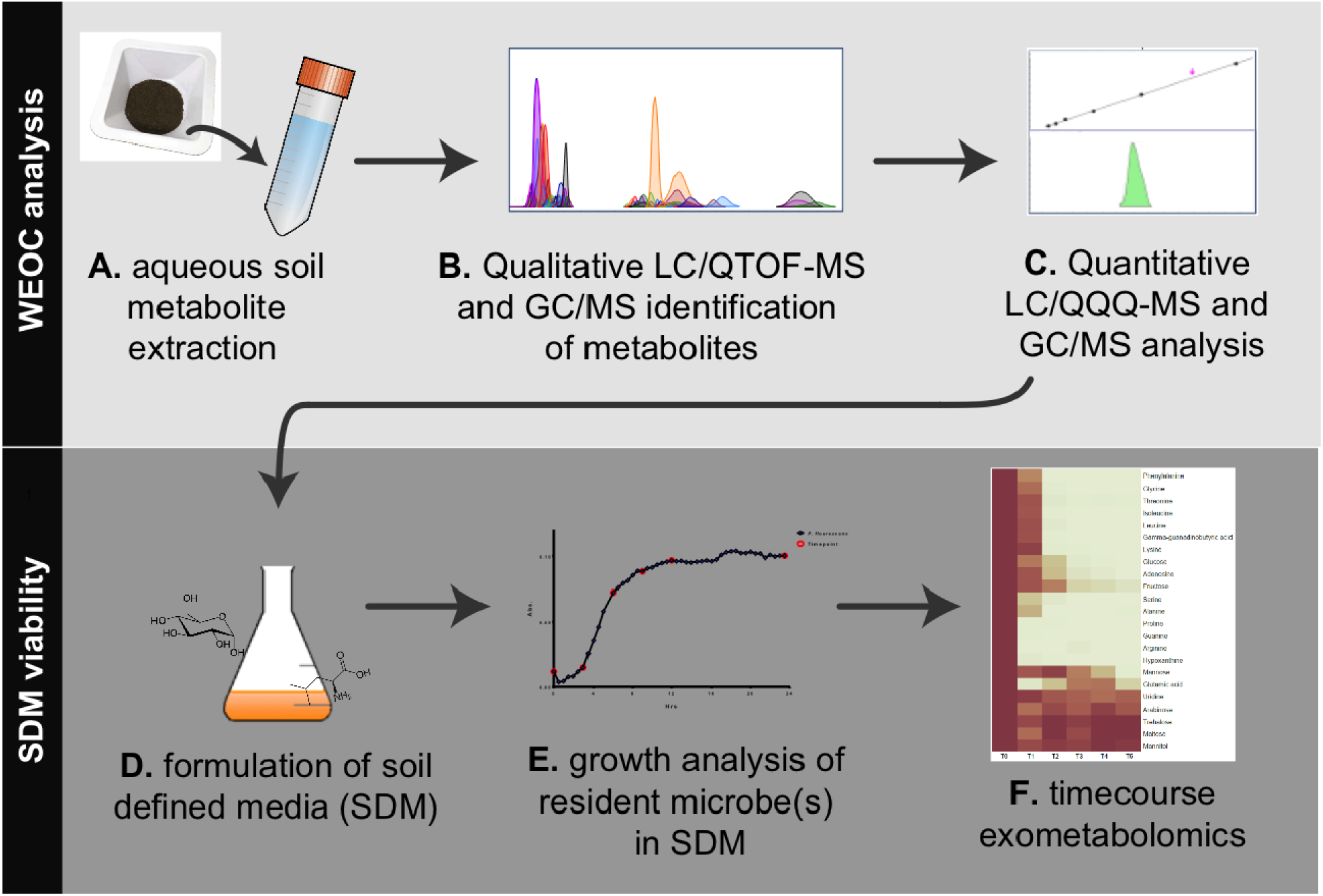
Workflow overview for the analysis of water extractable organic carbon (WEOC) and soil defined media (SDM). WEOC analysis is done by **(A)** extracting soil (sieved and fumigated) with water for 1 h, **(B)** acquiring scan and MS/MS data by LC/QTOF-MS and EI fragmentation data by GC/MS followed by WEOC metabolite identification and **(C)** quantitative analysis by LC/QQQ-MS and GC/MS using authentic standards. Based on these data, **(D)** SDM are formulated, (**E)** tested for viability using microbes isolated from the study site and **(F)** timecourse exometabolomics is performed to monitor substrate utilization by bacteria.

## 2 Methods and Materials

### 2.1 Chemicals

Glucose (CAS 50-99-7) was from Amresco (Solon, OH). LC/MS-grade methanol (CAS 67-56-1) and water were from J.T. Baker (Avantor Performance Materials, Center Valley, PA). N-methyl-N-trimethylsilytrifluoroacetamide (MSTFA) containing 1% trimethylchlorosilane (TMCS) was from Restek (Bellafonte, PA). Acetonitrile (CAS 75-05-8), adenosine (CAS 58-61-7), arabinose (CAS 28697-53-2), biotin (CAS 58-85-5), calcium chloride (CAS 10043-52-4), calcium pantothenate (CAS 137-08-6), cobalt nitrate (CAS 10026-22-9), d27-myristic acid (CAS 60658-41-5), ethylenediaminetetraacetic acid (EDTA; CAS 6381-92-6), folic acid (CAS 59-30-3), fructose (CAS 57-48-7), galactose (CAS 59-23-4), gamma-guanidinobutyric acid (CAS 463-00-3), guanine (CAS 73-40-5), hypoxanthine (CAS 68-94-0), iron sulfate (CAS 7782-63-0), magnesium sulfate (CAS 10034-99-8), maltitol (CAS 585-88-6), maltose (CAS 6363-53-7), manganese sulfate (CAS 10034-96-5), mannose (CAS 3458-28-4), mannitol (69-65-8), methoxyamine (CAS 593-56-6), nicotinic acid (CAS 59-67-6), p-aminobenzoic acid (CAS 150-13-0), monobasic potassium phosphate (CAS 7778-77-0), pyridine (CAS 110-86-1), pyridoxine hydrochloride (CAS 58-56-0), raffinose (CAS 17629-30-0), riboflavin (CAS 83-88-5), rutinose (CAS 90-74-4), sodium chloride (CAS 7647-14-5), sucrose (CAS 57-50-1), thiamine (CAS 67-03-8), thioctic acid (CAS 1077-28-7), trehalose dehydrate (CAS 6138-23-4), uridine (CAS 58-96-8), vitamin B12 (CAS 68-19-9), xylitol (CAS 87-99-0), xylobiose (CAS 6860-47-5), ^13^C-^15^ N-L-phenylalanine (CAS 878339-23-2) and amino acid kits #A6407 and A6282 were from Sigma-Aldrich (St. Louis, MO)

### 2.2 Aqueous soil extraction and total organic carbon analysis

Upper B-horizon soil was collected on September 24, 2013 from the background area of the ORFRC in Oak Ridge, Tennessee. Specifically, the collection site was located approximately 12 feet from groundwater well FW-300, in an area where grass, surface roots, and surface rocks were not abundantly present. To reduce the presence of debris and non-soil constituents from the soil sample, rocks and surface roots present on the surface, were physically removed prior to sampling. Surface soil was collected by shoveling soil to a depth of 12-inches into sterile whirl pack bags, which were immediately stored on blue ice. The soil at this location is an unconsolidated saprolite with organic rich clay soil that ranges from 0.5-3 m approximating the depth of the root zone. Other soil properties including total nitrogen, oxidizable organic matter, cation exchange capacity, pH, and particle size were determined by the UC Davis Analytical Laboratory (Davis, CA).

Aqueous extraction was performed as previously described (Swenson et al., 2015b). Briefly, soil was lyophilized to dryness and sieved to 2 mm prior to 24 h chloroform fumigation (Vance, 1987). Six grams of soil were extracted in 24 mL of water at 4°C for 1 h while shaking on an orbital shaker (Orbital-Genie, Scientific Industries, Bohemia, NY). Tubes were then centrifuged at 4°C for 5 min at 4000 rpm. Supernatants were collected in fresh 50 mL conical tubes and centrifuged again after which supernatants were filtered through 0.45 μm syringe filters (Pall Acrodisc Supor membrane). These final WEOC samples were lyophilized to dryness in a Labconco Freezone 2.5 (Labconco, Kansas City, MO) and resuspended in water at a final equivalent concentration of 2 g soil/mL. WEOC with and without further 1 kDa molecular weight cutoff (MWCO) filtration was analyzed for total organic carbon (TOC) by acidification and measurement of the non-purgeable organic carbon. WEOC samples were prepared as above, but at a final equivalent concentration of 1 g/mL (w/v of soil to water) and injected (50 μL) and run via an NPOC method using a Shimadzu (Kyoto, Japan) TOC-L series CSH/H-type TIC/TOC analyzer. Samples were individually mixed with a 1.5 % v/v of 1 M HCl for the conversion of inorganic carbon to CO_2_ followed by a 2.5 minute purging time and combustion of the remaining organic carbon at an oven temperature of 680°C. The carrier gas used was synthetic air at a flow rate of 80 mL/min. Values reported for TOC are in ppm (μg C per mL extract).

### 2.3 LC/Q-TOF Scan MS WEOC Profiling

One hundred microliters of the WEOC extract was lyophilized in triplicate to dryness and resuspended in 100 μL of methanol containing internal standards 3,6-dihydroxy-4-methylpyridazine, 4- (3,3-dimethyl-ureido)benzoic acid and 9-anthracene carboxylic acid at 25 μM. Samples were separated by hydrophilic interaction liquid chromatography (HILIC) on an Agilent 1290 UHPLC (Agilent Technologies, Santa Clara CA) using a Merck Sequant ZIC-pHILIC column (Merck KGaA, Darmstadt, Germany) of dimensions 150 x 2.1 mm and 4.6 μm particle size. Two microliters of sample were injected with the following mobile phase: A- 5mM ammonium acetate; B- 9:1 acetonitrile: 50 mM ammonium acetate. The following gradient was used with a flow rate of 0.4 mL/min: 0.0-1.5min, 100% B; 25.0 min, 50% B; 26.0-32.0 min, 35% B; 33.0-40.0 min, 100% B. The column temperature was set to 40°C and the autosampler was maintained at 4°C. Flow was directed to the electrospray ionization source of an Agilent 6550 quadrupole time-of-flight (Q-TOF) mass spectrometer with the following settings for both positive and negative polarity ionization modes; drying gas temperature and flow 275°C and 14 L/min; sheath gas temperature and flow were 275°C and 9 L/min, respectively; nebulizer pressure was 30 psi; capillary voltage was 3500 V and nozzle voltage 1000 V. Mass spectra were acquired from 50 to 1700 m/z at 4 spectra s^-1^.

### 2.4 LC/Q-TOF Tandem MS Analysis

Procedural blanks were used to generate exclusion lists of compounds and added to a data-dependent auto MS/MS method with a spectral scan range of 30-1200 amu and speed of 2 spectra sec^-1^. The quadrupole was set to a narrow isolation width of 1.3 amu and the collision cell fixed to 10, 20 and 40 eV. The auto MS/MS algorithm was restricted to a max of 4 precursor ions per MS/MS scan with a required precursor ion count threshold of 7,500 and a target of 50,000. Initial MS/MS spectra were then added within a retention time window of 0.25 min to the precursor exclusion list ensuring they were not selected for MS/MS in a subsequent run, a process that captured MS/MS spectra of low intensity precursor ions.

### 2.5 LC/QTOF-MS Data Analysis

MassHunter Qualitative Analysis v.B.07.00 SP1 was used to query acquired scan data against a custom LC method specific accurate mass and retention time metabolite database based on acquired authentic standard data. These initial metabolite identifications from scan MS data were augmented with tandem MS by searching at specific collision energies against spectral libraries such as METLIN (Smith et al., 2005), MassBank (Horai et al., 2010) and HMDB (Wishart et al., 2013). For metabolites only identified by LC/MS, the highest confidence identifications (ranked as “1” in Supplementary Table 1) required the unknown to be within 5 ppm m/z, 0.5 min retention time and to share dominant fragment ions with the standard. Lower confidence identifications matched standards in accurate mass and retention time but not fragmentation (or MS/MS was not obtained) and are ranked as a level “2” in Supplementary Table 1. For some metabolites, authentic standards were not available, but were identified based on matching fragmentation patterns with spectral libraries and are indicated by a level “3”.

### 2.6 GC/MS WEOC metabolite profiling

Sample extracts were dried and derivatized by oximation-silylation and data acquired by GC/MS as previously described (Swenson et al., 2015b). A C8-C30 fatty acid methyl ester ladder was added to each sample to enable orthogonal compound identification within a retention index window of +/- 20 with the electron ionization (EI) source of an Agilent 5977 GC/MS maintained at 70 eV for spectral library matching. Metabolite profiling data were deconvoluted using Unknowns Analysis v.B.07.00 from Agilent Technologies (Santa Clara, CA) followed by matching spectral fragmentation by quality score of 70 to the Agilent Fiehn GC/MS Metabolomics RTL Library (Kind et al., 2009). A set of sugar standards comprised of pentose, hexose, di-hexose and pentose alcohols (cellobiose, D-arabinose, D-arabinose, D-cellobiose, D-galactose, D-mannose, D-xylose, fructose, glucose, L-arabinose, maltitol, maltose, mannitol, mannose, raffinose, rutinose, sucrose, trehalose, xylitol and xylobiose) were used to resolve sugar pool composition for initial qualitative metabolite profiling and then prepared by single replicate at 1, 5, 10, 25 and 50 μM for determining sugar concentrations detected in the WEOC with d27-myristic acid used as an internal standard at a concentration of 25 μM (Supplementary Figure 1A). Quantitative data analysis was done using Agilent MassHunter Quantitative Analysis v.B.07.00. GC/MS identifications are provided in Supplementary Table 1.

### 2.7 LC/QQQ-MS for metabolite quantitation

For LC/MS-analyzed metabolites, quantitative assays were performed on an Agilent 6460 triple quadrupole (QQQ) MS (Agilent Technologies, Santa Clara CA) using the same LC, column and ESI source conditions as described for Q-TOF MS analysis. Authentic standards were prepared for amino acids, nucleobases and nucleosides for a subset of metabolites identified in the WEOC to generate compound specific transitions. Multiple reaction monitoring (MRM) transitions were scheduled by retention time segment as established using Agilent Optimizer v.B.07.00 and are provided in Supplementary Table 1. For the amino acid assay, a standard mix (Sigma-Aldrich amino acid standards kit, product#A6407, A6282) was diluted to concentrations of 0.1, 0.5, 1 and 10 μg/mL and run as single replicates (Supplementary Figure 1B). For the nucleotide/nucleobase assay, authentic standards were prepared at 1, 5, 10 and 25 μM concentrations in triplicate with ^13^C-phenylalanine used as an internal standard at 25 μM (Supplementary Figure 1C). Soil extracts were then run at a concentration of 2 g/ml (soil/water) for determining concentrations of these metabolites. Quantitative data analysis was performed using Agilent MassHunter Quantitative Analysis version B.07.00. For each metabolite, quantitative data (mg/mL for each metabolite) are converted to carbon ppm based on the percent weight carbon within each metabolite and are reported as mg kg^-1^ extracted soil.

### 2.8 Formulation of soil defined media and comparison of growth of 30 phylogenetically diverse ORFRC isolates

Soil defined medium 1 (SDM1) was prepared by adding 23 metabolites quantified from WEOC (Section 2.3) at the observed absolute concentrations, to a base media composed of 1x Wolfe’s mineral and 1x Wolfe’s vitamin solutions, potassium phosphate and ammonium chloride (Table 1). For soil defined medium 2 (SDM2), the number of metabolites was expanded to 46 to include additional observed, but not quantified WEOC compounds. These additional metabolites were added to the medium at the lowest concentration of a quantified metabolite within the same class (Supplemental Table 1). SDM (both at 1x and 10x concentrations) were compared to R2A medium at 1x concentration (Tecknova, Hollister CA) in their ability to support the growth of a broad range of ORFRC isolates. An existing collection of 30 phylogenetically diverse isolates from the ORFRC site were revived in liquid R2A from frozen glycerol stocks. DNA was obtained from overnight cultures to verify 16S rDNA based identifications. Aliquots were washed (3 times) prior to inoculation into 96-well microtitre plates. For each fresh medium, 20 uL of starter culture was added to 180 uL and plates were incubated under aerobic conditions for 110 h at 28°C, and shaken at 365 rpm. To facilitate growth monitoring, starting volumes of each isolate were normalized (200 uL total). Growth data were collected by measuring _OD600_ on a BioTek Eon Microplate Spectrophotometer at 15 min intervals.

**Table 1.** Metabolite formulation for soil defined medium 1 (SDM1). For each metabolite, concentrations are shown for soil WEOC (in mg/L), the amount added to SDM1 (mg/L) and the equivalent carbon ppm.

| Metabolite amendment | Formula | Molecular weight (g/mol) | Metabolite Class | WEOC mg/L | SDM1 mg/L | SDM1 C ppm |
| --- | --- | --- | --- | --- | --- | --- |
| Trehalose | C<sub>12</sub>H<sub>22</sub>O<sub>11</sub> | 342.3 | dihexose | 12.53 | 20 | 8.4 |
| Fructose | C<sub>6</sub>H<sub>12</sub>O<sub>6</sub> | 180.2 | hexose | 14.53 | 15 | 5.81 |
| Glucose | C<sub>6</sub>H<sub>12</sub>O<sub>6</sub> | 182.2 | hexose | 12.55 | 15 | 5.93 |
| Mannitol | C<sub>6</sub>H<sub>14</sub>O<sub>6</sub> | 182.2 | sugar alcohol | 8.17 | 15 | 5.93 |
| Maltose | C<sub>12</sub>H<sub>22</sub>O<sub>11</sub> | 342.3 | dihexose | 2.95 | 5 | 2.11 |
| Alanine | C<sub>3</sub>H<sub>7</sub>NO<sub>2</sub> | 89.1 | amino acid | 0.93 | 2 | 0.81 |
| Arabinose | C<sub>5</sub>H<sub>10</sub>O<sub>5</sub> | 150.1 | pentose | 0.52 | 2 | 0.8 |
| Leucine | C<sub>6</sub>H<sub>13</sub>NO<sub>2</sub> | 131.2 | amino acid | 0.7 | 1 | 0.55 |
| Mannose | C<sub>6</sub>H<sub>12</sub>O<sub>6</sub> | 180.2 | hexose | 0.54 | 1 | 0.4 |
| Isoleucine | C<sub>6</sub>H<sub>13</sub>NO<sub>2</sub> | 131.2 | amino acid | 0.46 | 1 | 0.55 |
| Arginine | C<sub>6</sub>H<sub>14</sub>N<sub>4</sub>O<sub>2</sub> | 174.2 | amino acid | 0.22 | 0.5 | 0.21 |
| Proline | C<sub>5</sub>H<sub>9</sub>NO<sub>2</sub> | 115.1 | amino acid | 0.2 | 0.5 | 0.26 |
| Threonine | C<sub>4</sub>H<sub>9</sub>NO<sub>3</sub> | 119.1 | amino acid | 0.2 | 0.5 | 0.2 |
| Lysine | C<sub>6</sub>H<sub>14</sub>N<sub>2</sub>O<sub>2</sub> | 146.2 | amino acid | 0.2 | 0.5 | 0.25 |
| Phenylalanine | C<sub>9</sub>H<sub>11</sub>NO<sub>2</sub> | 165.2 | amino acid | 0.19 | 0.5 | 0.33 |
| Glycine | C<sub>2</sub>H<sub>5</sub>NO<sub>2</sub> | 75.1 | amino acid | 0.17 | 0.5 | 0.16 |
| Uridine | C<sub>9</sub>H<sub>12</sub>N<sub>2</sub>O<sub>6</sub> | 244.2 | nucleoside | 0.16 | 0.5 | 0.22 |
| Glutamate | C<sub>5</sub>H<sub>9</sub>NO<sub>4</sub> | 147.1 | amino acid | 0.11 | 0.1 | 0.04 |
| Serine | C<sub>3</sub>H<sub>7</sub>NO<sub>3</sub> | 105.1 | amino acid | 0.09 | 0.1 | 0.03 |
| Adenosine | C<sub>10</sub>H<sub>13</sub>N<sub>5</sub>O<sub>4</sub> | 267.2 | nucleoside | 0.08 | 0.1 | 0.04 |
| Gamma-guanidino-butyric acid | C<sub>5</sub>H<sub>11</sub>N<sub>3</sub>O<sub>2</sub> | 145.2 | amino acid derivative | 0.05 | 0.1 | 0.04 |
| Hypoxanthine | C<sub>5</sub>H<sub>4</sub>N<sub>4</sub>O | 136.1 | purine | 0.04 | 0.1 | 0.04 |
| Guanine | C<sub>5</sub>H<sub>5</sub>N<sub>5</sub>O | 151.1 | nucleobase | 0.004 | 0.1 | 0.04 |
| **TOTAL** |  |  |  | **55.59** | **81.1** | **33.15** |

### 2.9 Time series exometabolomics analysis with *Pseudomonas sp.* FW300-N2E2

Using SDM1-10x liquid medium, triplicate 2 ml cultures (including media and water blank controls) of *Pseudomonas sp.* FW300-N2E2, a bacterial isolate from the ORFRC (16S rDNA sequencing closest to *Pseudomonas corrugata* strain P94- EF153018.1; Thorgersen et al. (2015)) were prepared in 24-well spectrophotometer plates using the inoculation technique described in 2.8. Time points were sampled early/mid log phase (3,6 hours) early/late stationary phase (9,12 hours) and finally at 24 h (Supplementary Figure 2). Culture fractions were removed for LC/MS and GC/MS analysis at 0.5 and 0.2 mL volumes, respectively, and centrifuged in a Savant centrifuge (Thermo Scientific, San Jose CA) for 5 min at 5000 rpm and the resultant supernatant filtered through 0.2 μm centrifugal filters (Pall Corporation, Port Washington NY) for 5 min at 10000 rpm. Filtrates were then frozen at -80 °C, lyophilized to dryness with LC/MS samples resuspended in 100 μL methanol containing 25 μM of the internal standards and analyzed as described in 2.3. GC/MS samples were derivatized and also analyzed as described in 2.6.

## 3 Results

### 3.1 Soil properties

The TOC of the WEOC was 129.1 ppm (or 129.1 mg C kg^-1^ soil). Other physical properties of the ORFRC soil were: total N of 0.08%, organic matter (oxidizable) content of 1.5%, cation exchange capacity of 14.5 meq/100 g, soil pH of 7.7 and particle size (% sand/silt/clay) of 29/54/17.

### 3.2 Integrating GC/MS with LC/MS to analyze soil WEOC composition and concentrations

In order to expand on GC/MS WEOC annotations and composition previously reported (Swenson et al., 2015b) this approach was complemented with HILIC LC/MS (Swenson et al., 2015a). Using both approaches here, a total of 96 water extractable soil metabolites were identified representing multiple classes of compounds such as amino acids, amino acid derivatives, mono- and di-carboxylic acids, nucleobases, nucleosides, osmolytes, sugars, sugar acids and sugar alcohols. Using LC/MS, 85 metabolites were identified (Supplementary Table 1) with 66 of these confirmed by analytical standards. Eighteen of the metabolites identified using LC/MS were also identified by GC/MS. In addition, there were 11 metabolites identified using GC/MS that were not detected by LC/MS. Sugars were best chromatographically resolved by GC/MS versus LC/MS. Using authentic standards, this sugar pool was found to be comprised of hexoses (glucose, fructose, arabinose, mannose), dihexoses (trehalose, maltose) and the sugar alcohol mannitol.

Quantitative methods were developed for 36 compounds representing the biochemical classes with the most abundant MS ion signal (amino acids, sugars, nucleobases and nucleosides). LC triple quadrupole MS was used for quantification of amino acids, nucleobases and nucleosides while sugars were quantified by GC/MS. Of these 36 quantified metabolites, 25 were within the quantifiable (linear) range (Supplementary Table 1). Converting the metabolite concentrations into carbon concentrations, these 25 metabolites together accounted for a total of 20.0 mg C kg^-1^ soil with sugars accounting for 89.2% of the quantified metabolites. Trehalose, glucose, mannitol, fructose and maltose each accounted for 5.3, 5.0, 3.2, 2.7,1.2 mg C kg^-1^ soil respectively (Supplementary Table 1). Of the remainder, 9.8% were amino acids and only 0.8% were nucleobases and nucleosides.

### 3.3 Molecular weight cutoff filtration and TOC quantitation of WEOC

Since the metabolomic methods used here would not be expected to detect all WEOC components, molecular weight cutoff filtration was used to help estimate the fraction of the WEOC pool quantified by MS. The TOC of the initial soil WEOC was 129.1 mg C kg^-1^ soil, which, after filtration through 1 kDa filters, was reduced to 53.3 mg C kg^-1^ soil. This reduction in TOC is consistent with the removal of biopolymers, colloids, trace lignin and cellulosic debris. Based on these TOC data, the 25 metabolites that were quantified from the WEOC, accounting for 20.0 mg C kg^-1^ soil, represented approximately 15.5% of the initial WEOC fraction and 37.5% of the <1 kDa filtered fraction.

### 3.4 Comparison of soil defined media and growth of bacterial isolates

Based on soil WEOC data, two soil defined media (SDM) were formulated. The first (SDM1) was based on 23 abundant (quantified) WEOC metabolites (Table 1). The second (SDM2) contained twice as many compounds (Supplementary Table 2) to determine if expanding the number of metabolites present in SDM supported additional growth of bacterial isolates. These additional metabolites (including many amino acids, nucleobases and nucleosides) were detected in the soil WEOC, but were not quantified. Because of this, they were added at the lowest concentration of a quantified metabolite within the same class. For both SDM1 and SDM2, added metabolites served as microbial carbon sources and were supplemented with 1x Wolfe’s vitamins and 1x Wolfe’s mineral solutions (Table 1).

A broad assessment was then performed to compare the viability of these two media, at two different concentrations, with 30 native ORFRC bacterial isolates (Figure 2). In order to benchmark the viability of these media, isolate growth was also compared to the isolation medium, R2A, which supported the growth of all 30 isolates. At 1x concentrations, SDM1 was able to support the growth of 11 isolates and at 10x SDM1 supported the growth of 13 isolates. While only five isolates grew on 1x SDM2, excitingly, 25 isolates grew on this medium at 10x concentration (Figure 2 and Supplementary Table 3).

**Figure 2.**
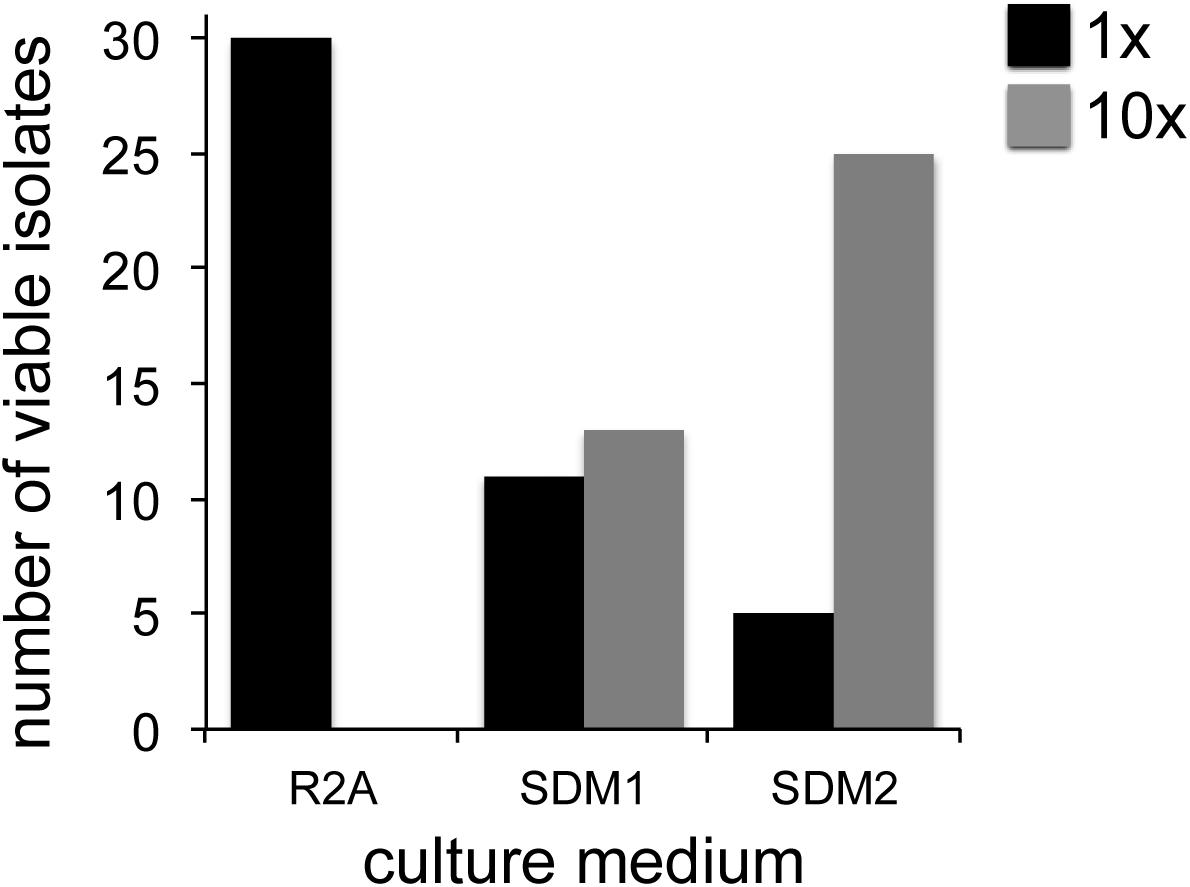
Isolate growth screen with R2A, SDM1 and SDM2. Each medium was tested (at 1x concentration for R2A and SDM and 10x for SDM) in its ability to support the growth of 30 phylogenetically diverse isolates from the ORFRC.

### 3.5 Isolate exometabolomics analysis using SDM1

Time-series exometabolomics studies can help delineate substrate preference of isolates as an additional measure of resource partitioning. Since SDM1 appeared to be a simple, yet viable medium for many isolates, this medium was selected to test its suitability for exometabolomics. *Pseudomonas sp.* FW300-N2E2 was grown in SDM1-5x and samples were collected at 3, 6, 9 and 12 h for targeted exometabolomics analysis by LC/QQQ-MS and GC/MS as described. Distinct patterns of substrate depletion were observed as shown in Figure 3. Compounds such as arginine, proline, glutamic acid, guanine and hypoxathine were already depleted by the first time point of 3 h. During later exponential growth phase (6 and 9 h), a second pool of compounds were depleted including alanine, isoleucine, gamma-guanadinobutyric acid, glycine, leucine, lysine, phenyalanine, serine and threonine. Interestingly, presumably high value substrates were not depleted until stationary phase with glucose detected up to 12 h and mannose and fructose up to the final 24 h sampling. Several metabolites persisted at near initial concentrations for the duration of the experiment including the abundant dihexoses trehalose and maltose, the hexose arabinose, the sugar alcohol mannitol and the nucleoside uridine.

**Figure 3.**
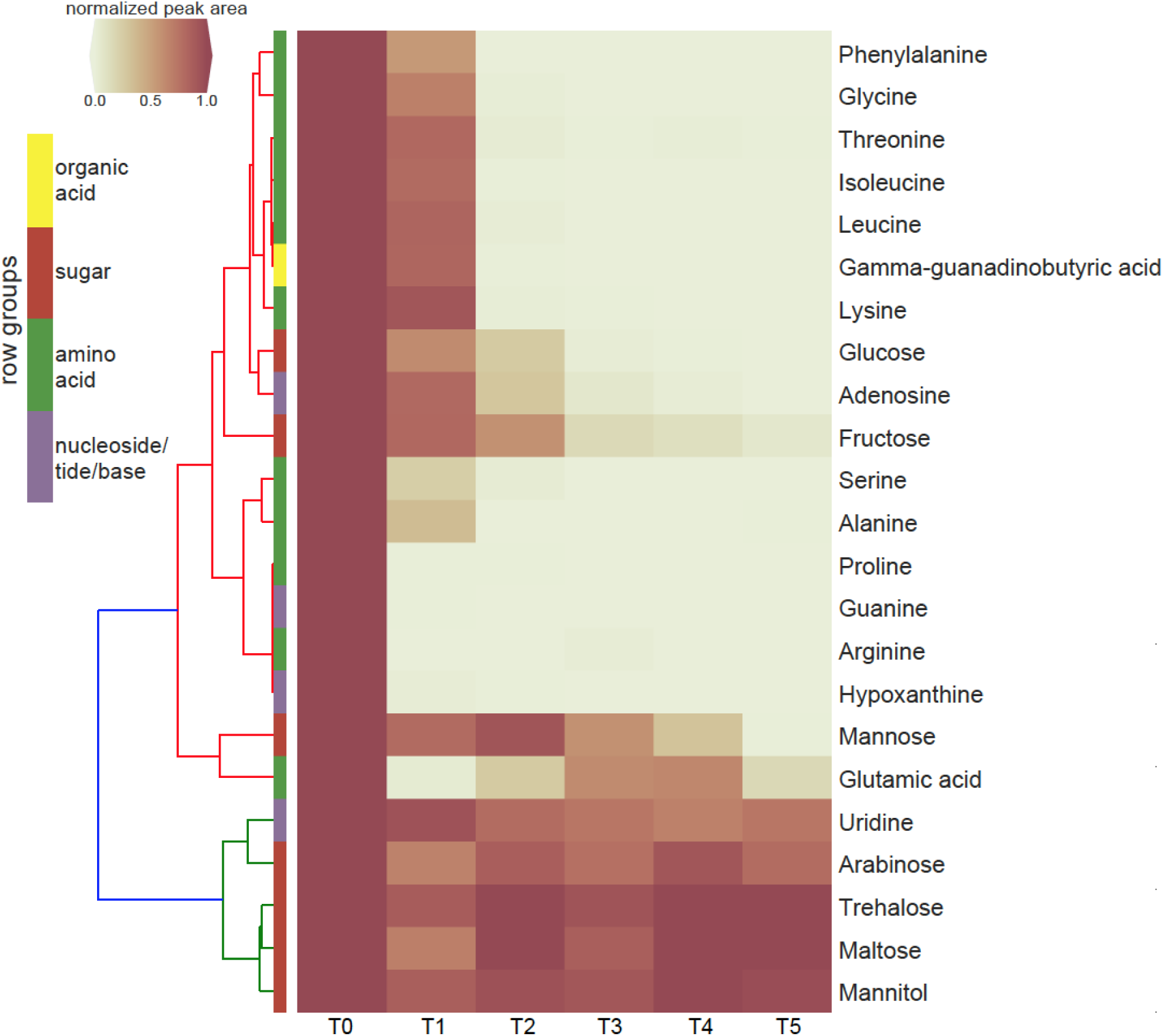
Clustering heatmap of normalized peak areas for SDM1 metabolites across timecourse sampling of *Pseudomonas sp.* FW300-N2E2 spent media. Levels are displayed in terms of relative ratio to initial concentration at time zero (T0) with T0-5 representing 0, 3, 6, 9, 12 and 24 h time points, respectively. Metabolite row groups are colored according to the metabolite class they belong to.

## 4 Discussion

### 4.1 Integrating qualitative GC/MS with LC/QTOF-MS to analyze WEOC composition

The objective of this study was to develop a viable and relevant defined media based on metabolomics analyses of an environment of interest to enable exometabolomic characterization of isolate resource partitioning. Here we combined two metabolomics approaches, GC/MS (Swenson et al., 2015b) and HILIC LC/MS (Swenson et al., 2015a) for qualitative and quantitative analysis. In total, 96 metabolites were identified using a combination of authentic standards, MS/MS and spectral library matching (Supplementary Table 1).

While this work focused on a single soil for the development of a metabolomics workflow for defined media preparation, it is informative to see how our metabolites detected lie within the context of other soil metabolomics studies. Overall, the range of metabolite classes detected, including amino acids, amino acid derivatives, mono- and di-carboxylic acids, nucleobases, nucleosides, osmolytes, sugars, sugar acids and sugar alcohols are consistent across many soil studies. The detection of trehalose and other compatible solutes (betaine and proline betaine) as well as additional quaternary ammonium compounds (acetylcarnitine, carnitine and choline) have been reported for other soils (Baran et al., 2013; Warren, 2014; Bouskill et al., 2016). Metabolites such as acetylcarnitine, citrulline, cytosine, gamma-aminobutyric acid and nicotinic acid are consistent with a key finding from Warren (2013) who showed that a diverse pool of nonpeptide organic N exists in soils.

The pool of organic acids in our WEOC samples was found to be diverse, composed of aliphatic di-carboxylic acids (malic, maleic, succinic) and aromatic carboxylic acids (benzoic, salicylic and shikimic). This is consistent with the observation of these low molecular weight carboxylic acids in many soils as reviewed by Strobel (2001) and the ability of aqueous shaking extraction to desorb higher concentrations of these metabolites than solution displacement techniques such as lysimeter collection or soil centrifugation. Our samples, which originated from an upper B horizon subsoil, may be associated with limited amounts of aliphatic mono-carboxylic acids (not detected in our WEOC extracts) potentially due to rapid microbial turnover of these compounds. Furthermore, the di-carboxylic and aromatic carboxylic acids that were detected here may be associated with a limited nutrient pool *in situ* that became desorbed during the aqueous shaking extraction (Strobel, 2001).

### 4.2. Analyzing the TOC content of soil WEOC

The TOC level observed in our soil WEOC (129.1 mg C kg^-1^ soil) was found to be consistent with many previous reports on soil DOC, noting that differences in soil types and extraction methods may dramatically affect these values. With this caveat, our TOC level is close to the 146 ppm reported for a soil (leachate) solution from a eutric cambisol grassland (Jones and Willett (2006). However, this study displays the affect of extraction techniques on these values by reporting lower TOC values (55-70 mg kg^-1^ soil) for the same soil following aqueous shaking (rather than leachate collection). Another study that is consistent with our TOC results focused on evaluating dissolved organic matter dynamics in Greek vineyard soils (Christou et al., 2005). They report drastic seasonal variation of DOC levels in soil solutions ranging from approximately 100-400 ppm over a 12 month period (Christou et al., 2005) and displayed substantial differences between topsoil and subsoil with WEOC levels decreasing from 89 to 58 mg C kg^-1^ soil (Christou et al., 2006).

Since soluble polymers and small particles presumably account for a large fraction of the WEOC, we wanted to determine the fraction of this total carbon pool accounted for by the 25 quantified metabolites. The TOC level of the soil WEOC was found to be reduced by more than half (129.1 ppm to 53.3 ppm) following 1 kDa filtration. Since microbes are limited in their ability to directly uptake macromolecules and are dependent on extracellular deconstruction followed by transporting the resulting metabolites, these molecules smaller than 1 kDa likely represent the most directly accessible fraction of the WEOC for microbes. This fraction is most relevant to our metabolomics methods given that that one operational definition of metabolites is molecules less than <1 kDa (Holmes et al., 2008).

### 4.3. Characterizing the LMWOS fraction of WEOC by quantitative LC/QQQ-MS and GC/MS analysis

Based on the molar concentrations of the 25 quantified metabolites in the WEOC (consisting mostly of carbohydrates and amino acids) and the number of carbons in each metabolite, we determined that these quantified metabolites accounted for 20.0 mg C kg^-1^ in the soil sample. This represents 15.5% of the WEOC and 37.5% of the < 1 kDa metabolite pool, indicating that even using two highly-sensitive analytical approaches, the majority of soil metabolites remain rare, unidentified or undetected. Numerous other studies support this assertion of the many unannotated chemical features present in soil LMWOS (Ohno, et al., 2010; Warren 2013; Baran et al., 2015) highlighting the value of using defined metabolite mixtures for exometabolomic characterization of microbial resource use.

The soil metabolites that we detected have a very uneven abundance distribution. Specifically, of the quantified metabolites, sugars represented approximately 89.2% of the quantified metabolite TOC pool, with a single metabolite, trehalose accounting for 5.3 mg C kg^-1^ soil (29.7% of the quantified sugar TOC pool and 9.9% of the < 1 kDa WEOC). Of the amino acids, only alanine, valine, leucine and isoleucine were at levels above 200 μg C kg^-1^ soil while all nucleosides and nucleobases except for uridine were at trace levels at or below 50 μg C kg^-1^ soil. While this type of metabolite distribution is specific to the ORFRC soil, we can compare these values with previous reports. Where we found total quantified amino acids to be 4.08 mg kg^-1^ soil, Fischer et al. (2007) examined the composition of Haplic Luvisol soil leachate and reports an amino acid content of 281.1 μg kg^-1^ soil, considerably lower than ours, though with a similar ranked composition (*e.g.* alanine, leucine and isoleucine ranked among the highest)(Fischer et al., 2007). We found the ratio of carbohydrates to amino acids to be 10.8 (g/g; more carbohydrates), which is consistent with Hertenberger et al (2002) who reports ratios ranging from 6.4-17.4, but Fisher et al. (2007) reports 0.4 (more amino acids). While these differences may simply be due to actual differences between soils, bias introduced during extraction is likely another important factor. For example extraction techniques range from mild soil leaching used by Fisher et al (Fischer et al., 2007) to the fumigation-aqueous extraction (this study) to a more aggressive acetone-water extraction used by Hertenberger et al. (2002).

### 4.4. Preparation and evaluation of defined media based on soil WEOC composition

SDM1, which contained 23 abundant WEOC metabolites, was found to support the growth of 13 out of the 30 isolates tested. Doubling the number of metabolites in the media (SDM2) dramatically increased the number of isolates that grew (25 out of 30) demonstrating similar viability to the widely used rich R2A media. Interestingly however, at reduced concentrations (1x), SDM1 supported growth of more isolates compared to SDM2. It could be that at 1x concentrations, SDM1 has a more optimal carbon concentration or C:N than SDM2, but at 10x concentrations, metabolite-specific transporters become active, promoting the survival in the more rich SDM2 (Button, 1993). While, the medium R2A was still more viable than the both SDM, this is not surprising given that the bacterial isolates analyzed here were isolated using R2A. Furthermore, R2A is rich in amino acids and peptides while SDM1 and SDM2 have a high sugar content, indicating potential unique and developed substrate preferences of these bacteria.

The application of SDM1 to investigate the substrate preferences of *Pseudomonas sp.* FW300- N2E2 revealed an interesting pattern of substrate utilization. We observed rapid depletion of arginine, glutamate, proline and guanine and hypoxanthine by the 3 h timepoint associated with the onset of logarithmic growth followed by consumption of alanine, isoleucine, gamma-guanadinobutyric acid, glycine, leucine, lysine, phenyalanine, serine and threonine (depleted by 6 h) then finally depletion of sugars including glucose. Several sugars such as arabinose, maltose, mannitol and trehalose were not used at all suggesting that this microbe may lack a sufficient set of transporters to use these abundant resources, supporting the view that it is important to perform exometabolomics experiments on media relevant, whenever possible, to their native environment. The utilization profile of this *Pseudomonas sp.* is consistent with a previous report for *Pseudomonas aeruginosa* PAO1 in which growth on a complex tryptone medium resolved an initial preference for select amino acids including leucine, proline, serine and threonine while the carbohydrate utilization was very similar to our results, namely, while glucose was metabolized, maltose, mannitol, mannose and trehalose were not (Frimmersdorf, Horatzek et al. 2010). Since, *Pseudomonas sp.* FW300-N2E2 was isolated using R2A, which is rich in many of the preferentially consumed compounds (such as amino acids) perhaps it is not surprising that it preferentially uses resources that, while relatively rare in the soil environment, are abundant in the isolation medium. We anticipate that media prepared based on soil metabolite analyses may have additional utility for isolating organisms that utilize the major measurable carbon sources within those environments.

The success of SDM demonstrates the ability to use soil metabolomics to develop defined media based on metabolites known to be abundant in soils and at concentrations relevant to the native environment. This also indicates the potential, in agreement with previous approaches, of using single or complex amendment of minimal media to isolate previously unculturable soil bacteria (Sait et al., 2002; Joseph et al., 2003). The approach used here is especially important for exometabolite profiling studies. Specifically, microbial transport and regulatory systems, particularly catabolite repression, are responsive to the absolute concentrations of metabolites in the environment. Thus, evaluating substrate utilization in defined media relevant to their habitat can greatly improve exometabolomic studies. There are a number of challenges associated with creating media that reflect microbial habitats for laboratory experimentation must be considered. Specifically, bulk analysis of soil no doubt creates averaged metabolite compositions unlike any particular niche within the soil. In addition, as we have shown here, many metabolites cannot be identified or go undetected. Complementing mass spectrometry with other types of spectroscopy, such as NMR, may help address this. Finally, due to the extreme heterogeneity between and within soil ecosystems, our particular findings on soil metabolite abundances should not be generalized except for the dozen or so metabolites that we have found to be consistent with other reports.

### 4.5. Implications for other environments

These methods, using exometabolomics analysis of environmental samples to prepare defined media for the study of organisms from that environment, are likely applicable to a diversity of environments. Extension of this approach to diverse soil types offers both the exciting possibility of helping connect soil metabolite composition to soil microbial community composition and development of generalizable defined soil media. While these defined media were focused on carbon, an important extension would be to also account for other critical elements especially organic nitrogen and phosphorus. The great advantage of these defined media is that they enable quantitative analysis of resource utilization by soil microorganisms.

One exciting implication of this analysis is that rare metabolites that were not analyzed may together account for a significant portion of the carbon (WEOC) in this sample, similar to how rare microbes collectively account for a large portion of microbiomes. Synthesis of this view with previous results showing that microbes from this soil (and biological soil crusts) utilize largely non-overlapping metabolites suggests there is coupling between microbial diversity and soil metabolite diversity. These results showing the unevenness of water soluble metabolites may further support the traditional view of copiotrophic and oligotrophic organisms. Specifically, there are a low-diversity of organisms competing for the abundant resources (copiotrophs) and a high diversity of rare organisms using low-abundance substrates (Upton and Nedwell, 1989; Konopka et al., 1998). It should be emphasized that this is highly speculative and studies of multiple soil samples and multiple sites would be required to support generalization to both this particular site and to soils in general.

## 5. Conclusion

This study used soil metabolomic analyses to characterize the low molecular weight organic matter composition to formulate defined media intended to approximate the qualitative and quantitative composition of microbe bioavailable carbon from a specific study site. Composition and carbon concentrations were found to align well with related studies with these soil metabolites having a very uneven quantitative distribution (e.g. trehalose accounting for 9.9% of the <1 kDa WEOC fraction), analogous to the uneven soil microbial community structure. The defined media that were synthesized to reflect the soil WEOC composition were found to support the growth of 13 or 25 (for SDM1 and SDM2, respectively) out of 30 phylogenetically diverse isolates. A detailed study of a single isolate, *Pseudomonas sp.* FW300-N2E2 showed that this isolate rapidly depleted guanine, serine, leucine and hypoxanthine while several metabolites including the most abundant disaccharides were not utilized from SDM1. We anticipate that this approach of preparing environmentally relevant defined media will be applicable to diverse environments to enable more ecologically relevant isolation and examination of microbial substrate utilization.

## Acknowledgements

The authors gratefully acknowledge funding from Ecosystems and Networks Integrated with Genes and Molecular Assemblies (http://enigma.lbl.gov), a Scientific Focus Area Program at Lawrence Berkeley National Laboratory for the U.S. Department of Energy, Office of Science, Office of Biological and Environmental Research under contract number DE-AC02-05CH11231.

